# Guided Diffusion for molecular generation with interaction prompt

**DOI:** 10.1101/2023.09.11.557141

**Authors:** Peng Wu, Huabin Du, Yingchao Yan, Tzong-Yi Lee, Chen Bai, Song Wu

## Abstract

Molecular generative models have demonstrated their potential in designing molecules from scratch with high binding affinities in a pre-determined protein pocket and could be combined with traditional structural based drug design strategy. However, the generative processes of such models are random and the atomic interaction information between ligand and protein are ignored. On the other hand, the ligand has high propensity to bind with residues called hotspots. Hotspot residues contribute to the majority of the binding free energies and have been recognized as appealing targets for designed molecules. In this work, we develop an interaction prompt guided diffusion model-InterDiff to deal with the challenges. Four kinds of atomic interactions are involved in our model and represented as learnable vector embeddings. These embeddings serve as conditions for each residue to guide the molecular generative process. Comprehensive in-silico experiments evince that our model could generate molecules with desired ligand-protein interactions in a guidable way. Furthermore, we validate InterDiff on two realistic protein-based therapeutic agents. Results show that InterDiff could generate molecules with better or similar binding mode compared to known targeted drugs.

## Introduction

Structure based drug discovery (SBDD) strives for design molecules that can bind to a target protein with high binding affinity and specificity, which acts as a critical approach in contemporary biopharmaceutical research[1]. However, SBDD remains a challenge owing to the massive chemical space. It is estimated that the number of “drug-like” molecules range from 10^20^-10^60^ considering the oral bioavailability and Lipinski’s rule-of-five[2]. Traditional *in silico* SBDD methods, such as virtual screening are computationally costly due to the large feasible chemical space and could not find novel drugs. In recent years, molecular generative models have emerged as a promising technique in drug discovery and enabled *de novo* molecular generation. Earlier methods relied on 1D (SMILES strings and SELFIES strings) or 2D (graphs) molecular representations[3-6], and are able to generate diverse and novel molecules. Nevertheless, these models ignore 3D spatial information of molecules and the protein pocket environment, which is essential for molecular properties and protein binding affinity. Consequently, 3D structure-based generative methods have gained lots of attention due to their capability of designing molecules that bind to a specific protein pocket.

Recently, various models have been proposed for 3D structure-based molecular generation, including variational autoencoders (VAEs), flow-based models, autoregressive models and diffusion models[7-14]. In [7], Matthew et al. represent molecules as density grids and a conditional VAE is used to generate new atomic density grids, then atom fitting and bond inference are applied to obtain novel molecules. Although they achieve remarkable results in generating diverse molecules, as pointed in [9, 10, 13], inferencing molecules from density grids is a nontrivial task and irregularities are presented in the generated molecules. Besides, the model is not equivariant and hard to scale to large protein systems. Peng et al. utilize autoregressive model to generate molecules atom by atom in protein pocket[14], but the generation process is inefficient and the deviations are accumulated as a result of the sequential generation. For instance, if the first several atoms are placed at improper positions, this will incur bias in subsequent generation process. On the contrary, diffusion-based models sample atom types and coordinates simultaneously in the light of protein context. Concretely, diffusion model defines a noise schedule and add noise to the molecular geometry in forward process. In the backward (generation) process, the model learns to reverse the noise process to recover the true molecular geometry. There is no mismatch between the training and generating process in diffusion models. Further, geometric symmetries in molecular system like rotation, translation and reflection are respected in diffusion model to improve the generalization ability.

Nonetheless, diffusion models still face one limitation in real scenarios. Essentially, diffusion models pertain to a model class named score-based generative models[15]. Another member in score-based generative models is score matching, which estimates the score of data at different noise scales and samples by gradually decreasing noise levels[16, 17]. As Song et al. pointed, when the number of noise scales go to infinity, score-based generative models can be regarded as a stochastic differential equation (SDE)[15]. While sampling from the SDE, there exists a corresponding ordinary differential equation (ODE) sharing the same marginal probability densities. On this account, the diversity of generated molecules would be decreased[12]. Additionally, the binding mode of proteins with ligand are vital for understanding the biological processes. It has been found that only a fraction of residues in the pocket, called hot spots contribute most to the binding affinity[18, 19]. A mutation in hot spots can cause significant drop in binding affinity and even drive drug resistance in patients[20, 21]. In modern development of drugs, hot spots are crucial for rational drug design and one usually desire that the drug can form interactions with hot spots. However, current diffusion models ignore the protein-ligand interaction information and cannot customize the generated molecules.

Inspired by the fruitful progress of prompt-based learning in nature language processing, we develop a prompt-based diffusion model called InterDiff to tailor the binding mode of generated molecules in the protein pocket. Specifically, we introduce four kinds of learnable prompt embedding to indicate the interaction type of protein residues, including π-π interaction, cation-π interaction, hydrogen bond interaction and halogen bond interaction. We perform an empirical study on CrossDocked2020 dataset[22] and shows that InterDiff is able to generate molecules under prescribed interaction prompts with high probability. In addition, we validate our model on two well-known targets in neural systems and cancers respectively. Experiments implicate that InterDiff could generate molecules having similar binding mode with known targeted drugs. To the best of our knowledge, this is the first work that introducing interaction prompt in structured based drug design.

## Results

InterDiff leverages interaction prompts to guide diffusion model and design molecules, resembling the prefix-tuning which optimizes a small continuous task-specific vector (Figure 1 and methods part)[23]. As shown in Figure 1, a diffusion process is learned to transform the ligand data distribution into a normal distribution conditional on the protein environment and interactions. In the generative process, the protein pocket and interactions serve as the conditions to guide each denoising step and design molecules that satisfy certain interactions. To evaluate InterDiff, we first conduct the experiment on a benchmark dataset compared with three recently published methods on five general metrics. Furthermore, we evaluate the model performance in generating molecules with predefined interactions, which is the key characteristic of InterDiff. We also illustrate the potential of InterDiff in real scenarios. Two protein targets with targeted drugs are selected and InterDiff is assessed to design molecules with identical interactions.

**Figure 1.**
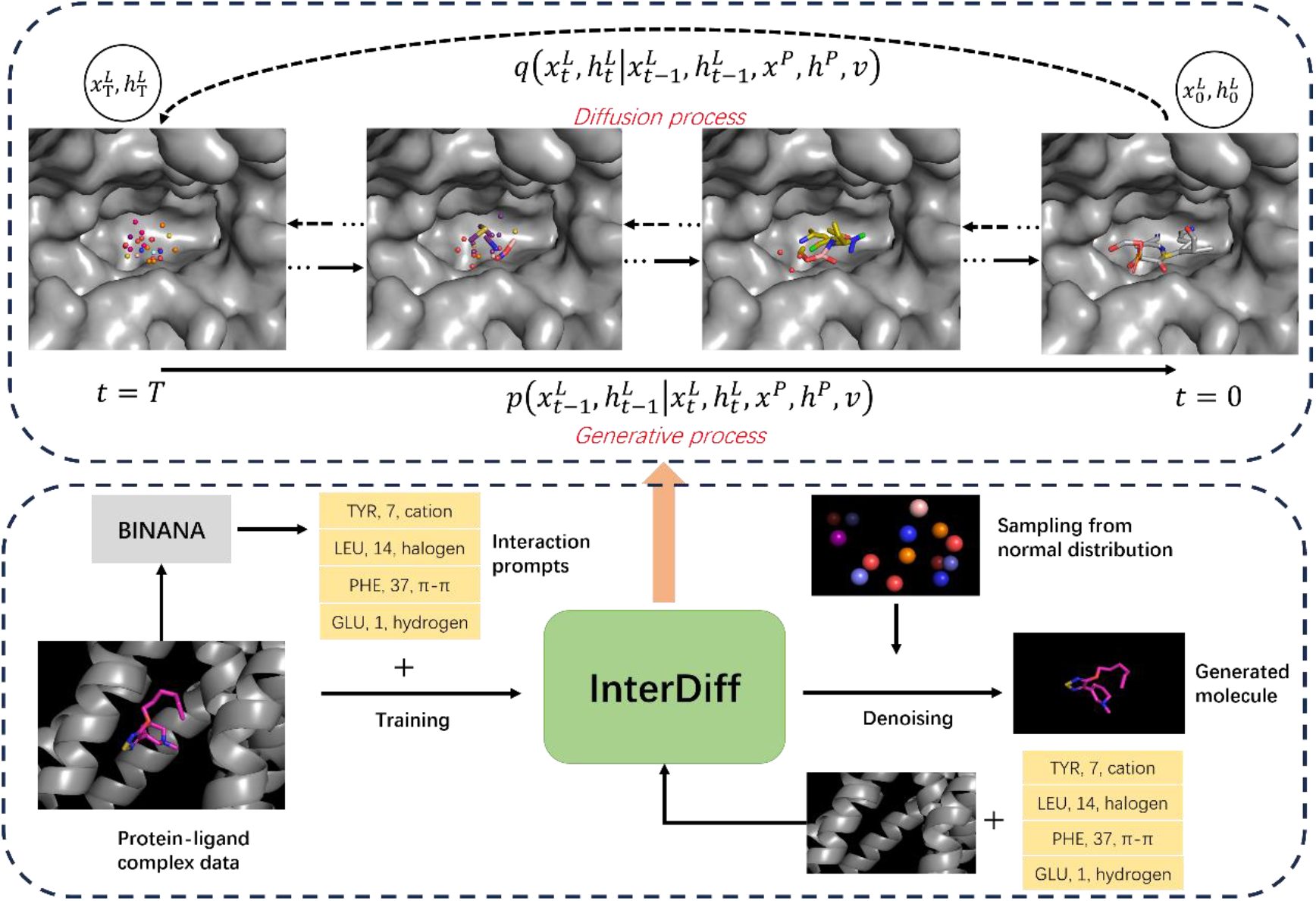
Workflow of InterDiff in a protein conditional generation. In diffusion process *q*, we simulate a progressively noised ligand point cloud (coordinates and atom types: *x*^*L*^, *h*^*L*^ under protein environment (*x*^*P*^, *h*^*P*^ over *T* timesteps. The interaction prompts *v* indicates the interaction types for each residue (‘*TYR, 7, cation’*: the 7^th^ residue is tyrosine with cation-π interaction, which are extracted by BINANA and are tunable in the diffusion process. In the generative process *p*, a neural network learns to recover data from a noise distribution conditioned on protein and prompts.

### Data

The crossDocked2020 was used to train and evaluate InterDiff[22]. We follow the same splitting and filtering criterion described in [10, 14], obtaining 100000 samples for training and 100 samples for testing. The protein-ligand interactions are detected by BINANA2[24], four kinds of interactions are adopted including cation-π, π-π, hydrogen and halogen interaction. We ignore the complexes that have residue detected with more than one interaction. Phenylalanine and Tyrosine were found to have four interactions (Figure S1) and hydrogen bond account for the vast majority of interactions (Figure S1, S2).

### Molecular structure and properties

Routinely, we assess the generated molecules from test set in five commonly used metrics: 1) **Vina Score**. A binding affinity indicator calculated by physical-based empirical scoring function. 2) **QED**. QED is a measurement of drug-likeness of molecules. 3) **SA**. SA (synthetic accessibility) measures the feasibility of synthesize molecule based on fragmental analysis in compound database. 4) **Lipinski**. We calculate the Lipinski score by quantifying how many rules are fulfilled in Lipinski’s rule of five. 5) **Diversity**. Diversity is computed by averaging the pairwise dissimilarity (one minus *Tanimoto* similarity) of generated molecules in each pocket.

Tab 1 displays the results of InterDiff and baseline methods. Overall, InterDiff outperforms baseline methods in the diversity and the other indicators are less ideal. We notice that the average number of atoms in generated molecules sampled by InterDiff is smaller than other methods, ranging from 2-4 atoms. Since a larger molecule tends to have a better docking score, this may account for the vina score for our method compared with TargetDiff. TargetDiff achieves best results in vina score and Pocket2Mol performs best in QED, SA and Lipinski. However, this also indicates that InterDiff and TargetDiff could generate novel molecules since QED and SA are calculated based on existing drug database. To evaluate the substructure of generated molecules, we count the percentage of different ring sizes. Results show that InterDiff and TargetDiff tend to produce a larger proportion of 7-membered ring while AR has more 3-membered rings. (Tab S1). This could partially explain the lower score in QED and SA of TargetDiff and InterDiff, since 7-membered ring rarely appears in common drugs[25]. Besides, we spot that the variance of vina score in InterDiff are much smaller than the other methods, as shown in Figure 2 and Table 1. In the generative process, we keep the interaction prompt of the protein residues the same (see methods part) as reference molecules in the test set and the number of atoms is sampled according to the distribution of pocket size and number of atoms in ligand (Figure S3). The interaction prompt for the residues restricts the fluctuating extent of binding pose of the generated molecules, resulting in a binding mode akin to the reference molecules. We will analyze the accuracy of InterDiff in generated molecules with defined interaction prompt. Considering that the vina score is calculated based on the binding pose of ligand, this could shed light on lower variance of vina score in InterDiff.

**Table 1:** Evaluation results from test set of CrossDocked 2020 dataset. The performance is re-evaluated for baseline methods Pocket2Mol, AR and TargetDiff.

| | Vina Score (↓) | QED (↑) | SA (↑) | Lipinski (↑) | Diversity (↑) |
| --- | --- | --- | --- | --- | --- |
| Test set | -6.871±2.32 | 0.476±0.20 | 0.728±0.14 | 4.340±1.14 | — |
| Pocket2Mol | -6.561±2.67 | **0.573**±0.16 | **0.756**±0.13 | **4.879**±0.42 | 0.731±0.12 |
| AR(3D-SBDD) | -6.592±2.08 | 0.507±0.19 | 0.634±0.14 | 4.723±0.65 | 0.698±0.10 |
| TargetDiff | **-7.163**±1.72 | 0.472±0.20 | 0.585±0.12 | 4.519±0.84 | 0.717±0.09 |
| InterDiff | -6.584±1.29 | 0.413±0.18 | 0.598±0.11 | 4.405±1.01 | **0.820**±0.04 |

**Figure 2.**
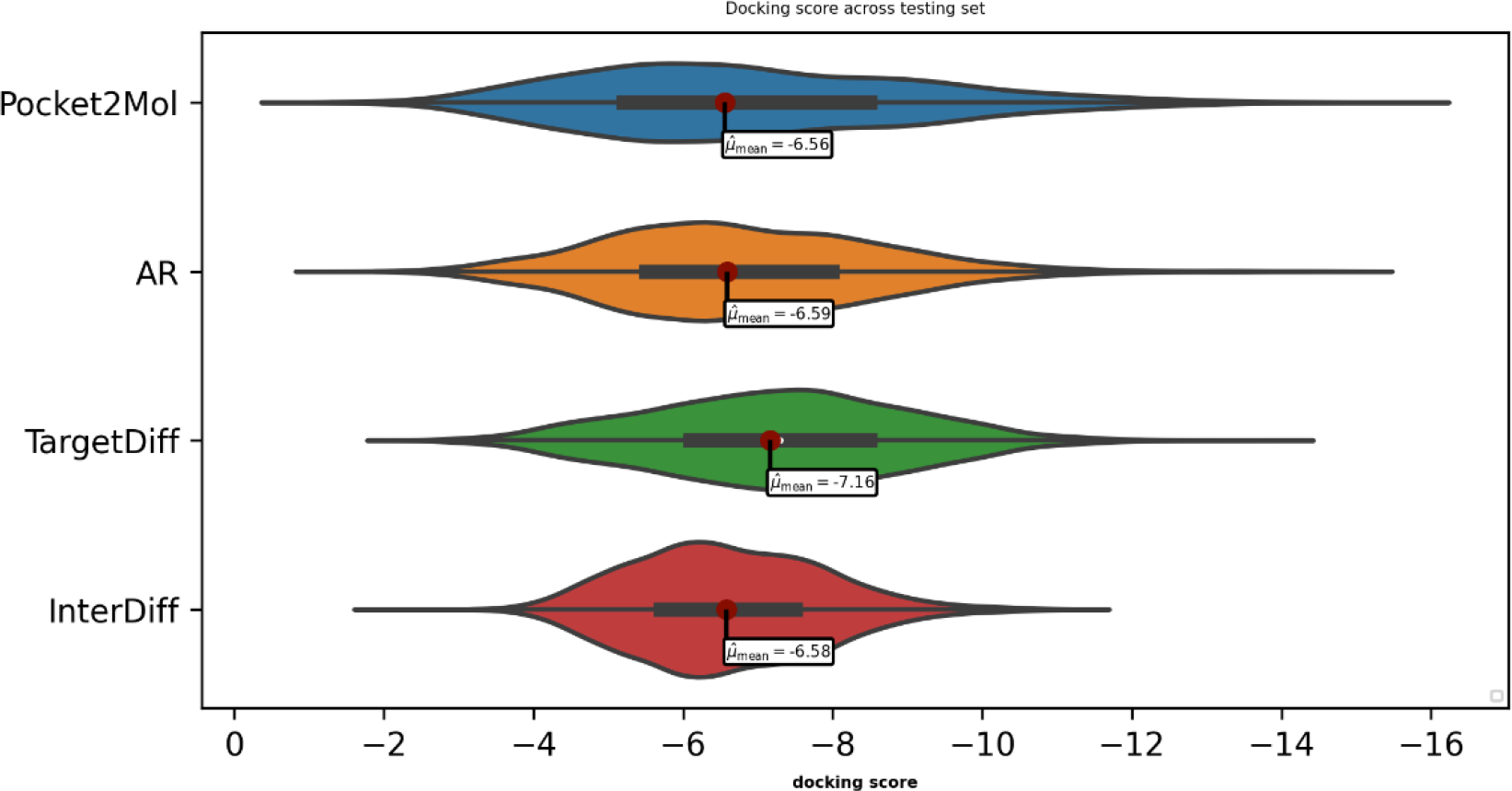
violin plot of distributions of docking score for InterDiff and baseline methods. InterDiff shows a limited range of scores compared to other methods.

We further evaluate the molecular chemical space distribution of generated molecules by 2D and 3D molecular fingerprints. The projection of Morgan Fingerprint and USRCAT (Ultrafast Shape Recognition with CREDO Atom types) features by UMAP (Uniform Manifold Approximation and Projection) are displayed in Figure S4 and S5. Morgan Fingerprint derives from 2D molecular representation and takes atom types and connectivity into account. USRCAT is a method that measures 3D shapes of molecules meanwhile considering the pharmacophoric features. Generally, InterDiff has similar distributions in space with TargetDiff and AR shows a dispersed distribution regarding USRCAT projection. Pocket2Mol behaves differently against others, which locates at the upper corner and may correspond to a region with more drug-like molecules. For the 2D Morgan Fingerprint, InterDiff has one dense region while other models have two or more than three dense regions. The interaction prompt may confine the diversities of structures presented in generated molecules, making them similar to the reference molecules. This is in line with the distribution of vina score, which has a narrow range comparing to other methods.

### Performance of InterDiff in design specific Interactions

To evaluate the capability of InterDiff in generating molecules with designated interactions, we re-generate molecules in the test set for 100 times and the number of atoms is identical to the reference molecule for each sample. Additionally, we excluded the test samples that did not detect any interaction and 99 samples are left for testing. After generation, the interactions were detected by BINANA2 using the conformers generated by InterDiff in protein pocket. We compute the accuracy of accomplishing designated interactions for generated molecules. As an illustration, if *[‘GLU’, 14, ‘Hydrogen’]* (fourteenth residue GLU with hydrogen bond) and *[‘TYR’, 39, ‘caption’]* are given as the condition for generating, and only *(‘TYR’, 39, ‘caption’)* is obtained in the generated molecular conformer, the accuracy would be 50%. The results are exhibited in Figure 3:

**Figure 3.**
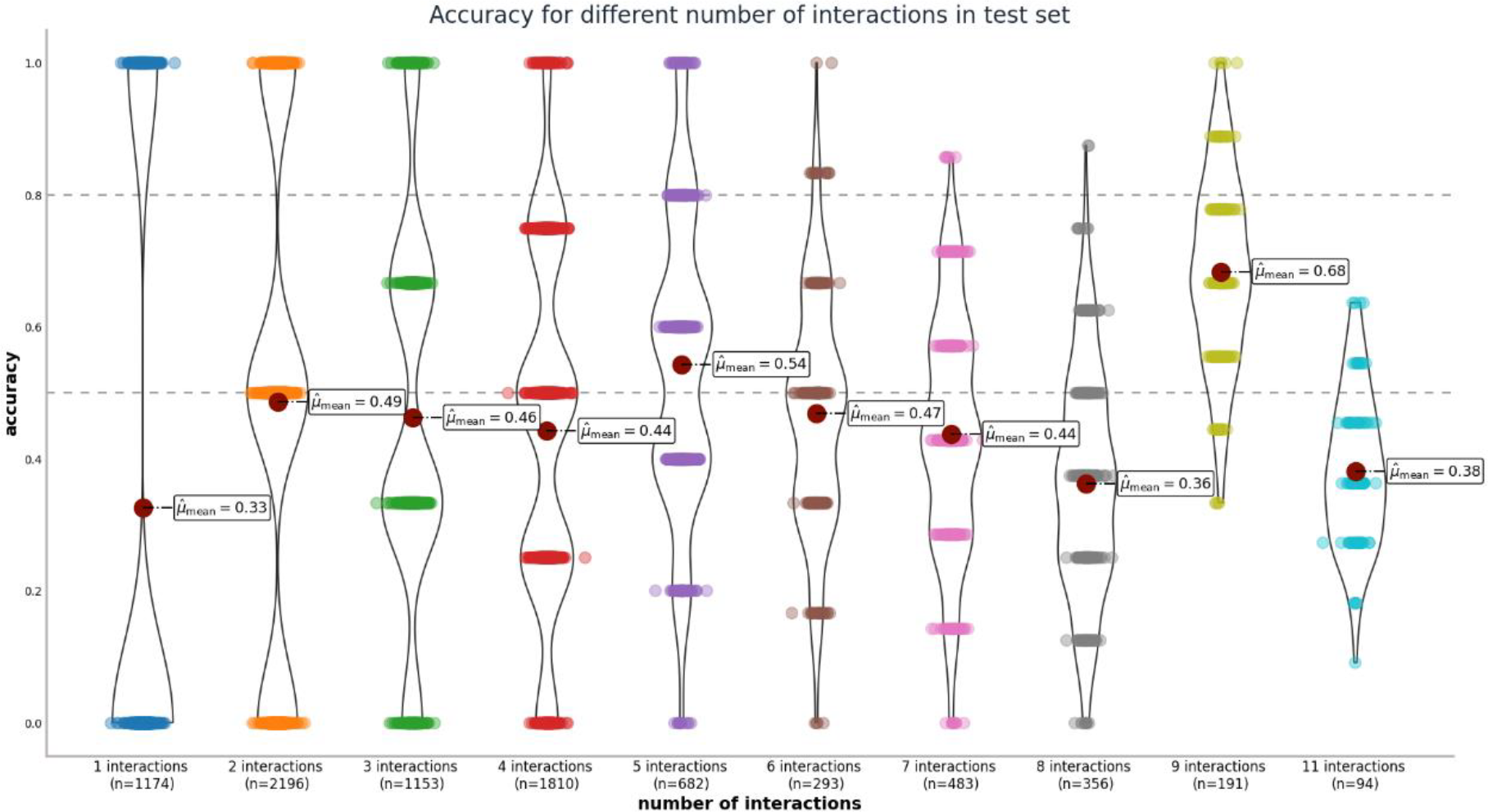
Accuracy of achieving designated interactions in generated molecules in test set. The samples are sorted by the number of interactions.

The number of interactions in the test set ranges from 1 to 11 except for the 10 interactions and we generate 100 samples for each protein pocket. Overall, InterDiff illustrates excellent performance in designing molecules under specific interaction prompts. We achieve the highest accuracy under 9 interactions (mean = 68%, n = 191) and the lowest accuracy under 1 interaction (mean = 33%, n = 1174). In the most difficult setting 11 interactions, InterDiff still reach an average accuracy of 38% in 94 generated molecules (6 molecules fail to reconstruct). Among the 94 generated molecules, 4 samples realize the highest accuracy with 7 interactions agreed with the reference. What’ s more, in 9 interaction cases, 5 molecules attain the same interaction types as the reference and 23 molecules are in accord with 8 interactions. To verify if the generated conformers agree with the pose after docking, we compare the raw conformers from InterDiff with the docking poses generated by QuickVina. We plot the resulting RMSD density distribution of 9 conformers (ordered by the docking score) generated by QuickVina (Figure S7). For the besting scoring pose, QuickVina agrees with 9% of generated conformers (RMSD below 2 angstrom), which is similar to the work that Schneuing et al. reported[9]. Additionally, we also observe a significant drop of the proportion of agreed molecules in the less confident poses (4% in the 9^th^ pose). This indicates InterDiff can generate molecular conformers approaching the steady binding pose. Furthermore, to estimate the ability of InterDiff in achieving disparate interactions, we assess the accuracy of four interactions in each sample and the results are shown in Figure S6.

Overall, InterDiff behaves distinctly in designing disparate interaction. The probability of hydrogen bonds being designed is the highest, followed by halogen bond. We note that InterDiff does not perform well in π-π interactions, which is understandable since the requirements of π-π interaction are more complicated than the others. Accordant with BINANA2’s criterion, three standardized aromatic residues (phenylalanine, tyrosine and histidine) are involved. The aromatic ring center distance between ligand and protein must be less than 4.4 angstroms and the ring atoms in could not deviate from planarity by more than 15 degrees. Last but not least, the angle of two normal vectors in the ring planes needs to within 30 degrees. Compared to other interactions, π-π interactions demand aromatic ring structure on the ligand and the ring has to be positioned in a proper manner. In addition, we noticed that the π-π interactions only comprise around 8 percent of the total interaction samples. On account of this, we speculate that the imbalanced interaction distribution may also impact the model performance in π-π interactions.

### Application of InterDiff in real scenarios

In this section, we investigate the potential of InterDiff in designing drugs when the binding mode of a reference molecule is available. We select two protein targets with different subtypes and use InterDiff to design molecules with similar binding mode as existing drugs. The first target is muscarinic acetylcholine receptor (mAChR), acting an important target in central nervous system diseases, for instance, Alzheimers’s disease and schizophrenia[26]. Xanomeline was developed as an agonist to mAChRs and studies have found that it has almost identical binding affinity to all mAChR subtypes (M1-M5), but stimulates them to appreciably different extent[27]. A recent study termed this phenomenon as “efficacy-driven selectivity” and the authors found that Xanomeline’s binding mode differs between inactive states and active states of mAChRs[28]. We use InterDiff to design molecules for M2 type mAChR in both inactive state and active state, conditioning on the binding mode of Xanomeline in two states. The second target is KRAS, commonly mutated in cancers and serving as a therapeutic targeting in various cancers, such as lung cancer, colorectal cancer and pancreatic cancer. Current inhibitors only target KRAS G12C mutants but the non-G12C mutants constitute the most in KRAS driven cancers. Recently, Kim et al. reported a non-covalent inhibitor BI-2865, which can bind to a wide range of KRAS altercations[29]. In like manner, we design molecules with InterDiff and take the binding mode of BI-2865 in KRAS G12C and another mutant G13D as references. We sample 300 molecules for each state or mutant of two targets and check the interactions after docking by QuickVina. The original interactions for two drugs are list in Table S3. Among the generated molecules, we successfully obtain molecules that have identical interactions as existing drugs, and we randomly select four molecules for illustration.

As demonstrated in Figure 4, we present the poses of generated molecules by docking in protein pockets together with the native targeted drugs. The poses of Xanomeline in mAChRs are also acquired by docking and the poses of BI-2865 in KRAS are acquired from cocrystal structure in PDB database. We can see that the docking pose of designed molecules overlap well with the reference drug. What’s more, InterDiff successfully generates similar interactions as the reference drugs in three of the four protein targets. For the last target (Figure 4d, KRAS wild type), three of the four interactions are consistent.

**Figure 4.**
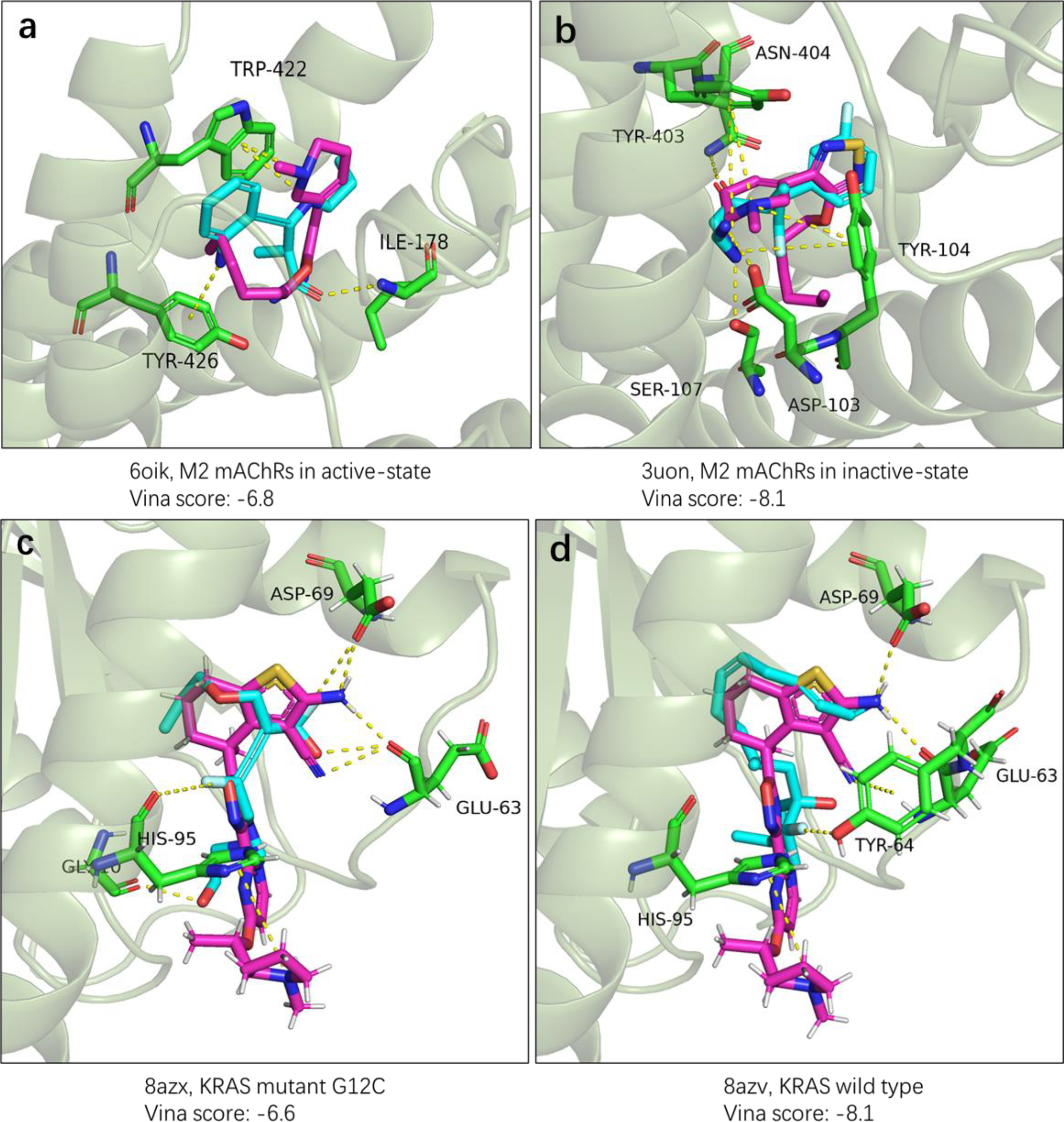
Pose of generated molecules and native drug in protein target. **a, b**: The illustration of Xanomeline and generated molecules in M2 mAChRs active state and inactive state. **c, d**: Generated molecules and BI-2865 in KRAS wild type and G12C mutant. The binding poses of four generated molecules and Xanomeline are obtained by QuickVina while the pose of BI-2865 are obtained from cocrystal structure. Residues that have interactions with molecules are colored with green. BI-2865 and Xanomeline are colored with purple and generated molecules are colored with blue. The structures of all molecules are available in Figure S8.

The docking pose of generated molecules and Xanomeline for M2 mAChRs in active and inactive state are illustrated in Figure 4a and 4b. For the active state (PDB:6oik), the original interaction is TRP-422 with interaction cation-π. InterDiff successfully realizes the same interaction and additionally introduces two new interactions, TYR-426 with cation-π and ILE-178 with hydrogen bond. For the inactive state (PDB:3uon), the primary interactions are cation-π in TYR-403 and TYR-104. InterDiff also reproduces the same interactions and three extra hydrogen bonds are formed in ASN-404, SER-107and ASP-103. In the second case, three interactions are discovered by BINANA2 in KRAS mutant G12C cocrystal structure (PDB:8azx), ASP-69 with hydrogen bond, GLU-63 with hydrogen bond and HIS-95 with cation-π. While in KRAS wild type (PDB:8azv), an additional interaction, TYR-64 with cation-π is found. InterDiff could discover molecules with the similar binding mode in KRAS mutant G12C (form an extra interaction GLY-10 with hydrogen, Figure 4c) and KRAS wild type except for the interaction HIS-95 with cation-π. Besides that, we notice that the BI-2865 has 5 ring structures, and there are no rings in molecules generated by InterDiff. Currently, InterDiff could not control the sub-structures in generating process and this could be a future direction.

### Fragment growing by inpainting

In this part, we investigate the potential of InterDiff in fragment-based drug design (FBDD). FBDD enables designing molecules conditioned on a potent substructure. It is very common that one may desire to optimize certain parts of a molecule while fix the molecular scaffold. To this end, we additionally train an unconditional diffusion model which learns the joint distribution of ligand atoms and protein atoms. The model structure and training process is identical to the conditional InterDiff except for the training objective (protein atoms are included in the loss function). To generate molecules under a given scaffold, we modify the sampling process by injecting the fixed context in the denoising steps and replacing the corresponding parts from the model. This technique is named inpainting and initially introduced in image imputation[30, 31]. Formally, in each denoising step, we do following operations:

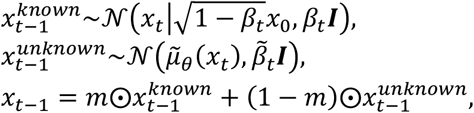

where 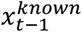 indicates the reference samples, 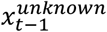indicates the samples from the model and *m* is a binary mask which signifies the fixed context. In the experiments, the pocket atoms and the native ligand are the 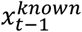and the denoising samples from the model are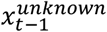. It is evident that by iterating this step during the sampling process the molecular scaffold can be preserved in the final generated molecules.

The same protein targets are used in experiment as the previous section and 100 molecules are sampled for each state of the two targets. We evaluate InterDiff in FBDD by removing the fragments (Figure 5, transparent parts) that have interactions with protein residues and keep the rest of the molecules as the fixed scaffolds. Four illustrative examples are shown in Figure 5, and we successfully design fragments with the same interactions as the native drug in M2 mAChRs based on the scaffold. For the KRAS mutant G12C (PDB:8azx) and KRAS wild type (PDB:8azv), one (ASP-69 with hydrogen) of three and two (ASP-69 with hydrogen, TYR-64 with cation-π) of four interactions are achieved. The docking poses are generated by the model and our method can inpaint new fragments with desired interactions around the fixed scaffolds. However, we also noticed that the accuracy of InterDiff in inpainting mode is lower than the pocket-conditional mode. The model has to estimate the positions of both protein and ligand atoms in the denoising steps and on the contrary, only ligand atoms are estimated in the pocket-conditional mode. The errors in estimating protein atom types and locations could influence the accuracy in the estimation of ligand atoms. In addition, in case PDB:6oik, we found that the new fragments are anchored in an alternative position on the 5-membered ring. It would be interesting to add the information of anchor point in the diffusion model and generate diverse molecules.

**Figure 5.**
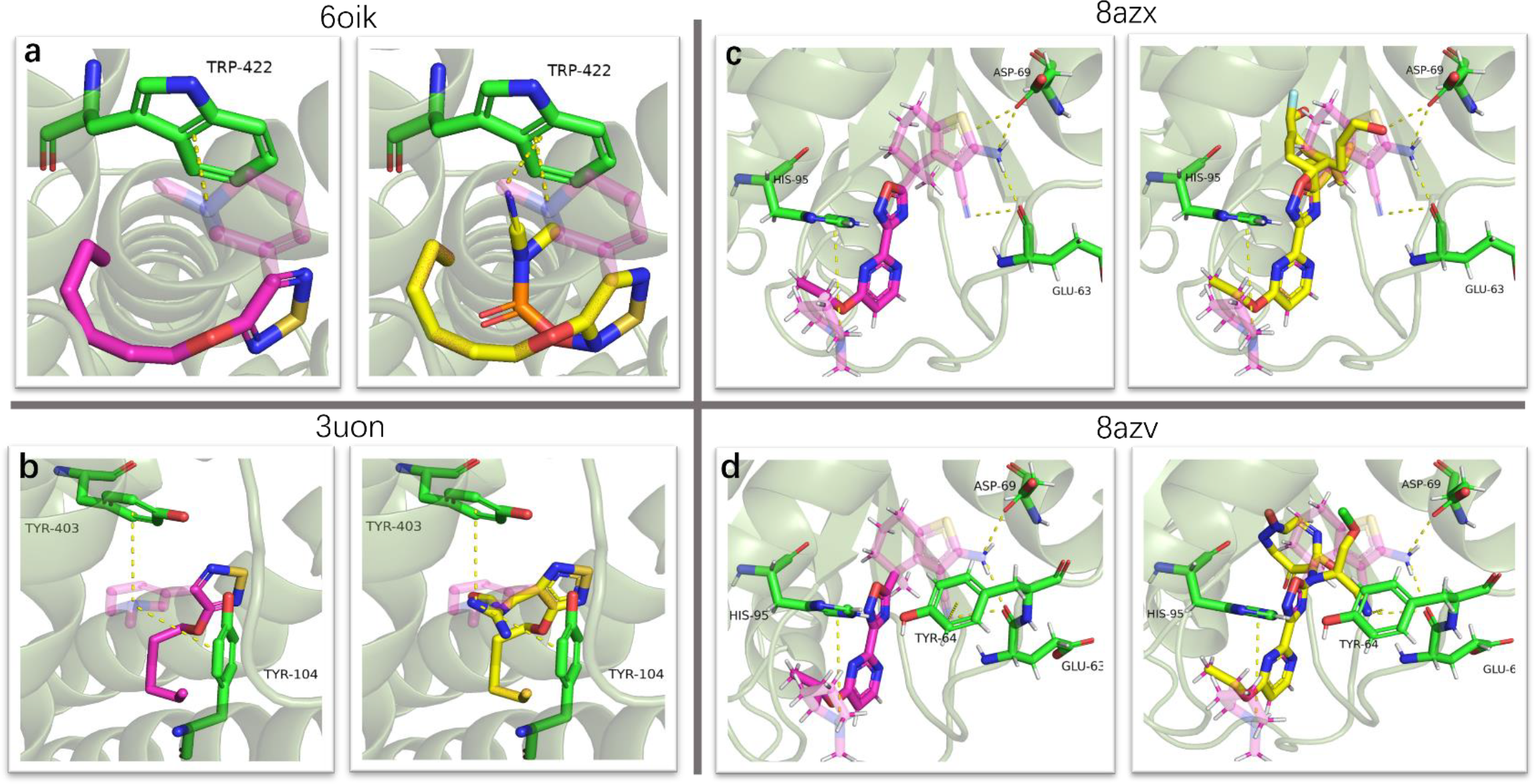
Pose of designed fragments with scaffold and native drug in the protein target. The transparent parts of native drug are fragments with atomic interactions. Residues that have interactions with molecules are colored with green. BI-2865 and Xanomeline are colored with purple and generated molecules are colored with yellow.

## Related work

### Diffusion model for molecular design

Diffusion models are a new kind of generative model inspired by diffusion process. Impressive progresses have been made in distinct generating task such as images, audios and even videos[32-34]. In molecular science, Hoogeboom et al. first proposed E(3) Equivariant Diffusion Model (EDM) for molecular generation which notably outperforms previous 3D generative methods[35]. Shortly after their work, Schneuing developed a diffusion model for structured based drug design named DiffSBDD, which is the first of its kind[9]. Two strategies are introduced under their framework, protein-conditional and ligand-inpainting generation. Specifically, ligand-inpainting method learns the joint distribution of protein-ligand complex, and new ligands are completed in inference stage. Experiments exhibit that both strategies can produce novel and drug-like ligands. In silico docking assessment also verify the potential in generating ligands with high binding affinity. Similar work were done in [13] and the difference lies in a dual diffusion was used to capture the local and global protein environment. In addition, Guan et al. presented a target-aware diffusion model. Unlike previous work that need to evaluate generated molecules through docking method like AutoDock, their model can estimate and rank the binding affinity of molecules. The authors raise a problem that the bond inference is implemented in a post-processing manner and irrational structures may appear in generated molecules, which is also pointed out in Schneuing’s work[9]. Huang et al. tackle this problem by setting distance threshold for covalent bond[12], but the bond distance can vary depending on the particular chemical structure. Alternatively, Wu et al. developed a diffusion model guided by a prior diffusion bridge[36], which can guarantee a desirable output. Specifically, AMBER inspired physical energy and statistical energy were incorporated as priors to guide training process. Another solution to this issue is integrating the chemical bonds into the diffusion process to enhance the quality of generated molecules[37].

### Prompt learning for molecular design

Prompt-based learning is initially a strategy to train large language models (LLMs), serving as an alternative to the fine-tuning paradigm so the LLMs can adapt to different tasks without re-training[38]. Afterward this technique was introduced to vision-language model and greatly improved the performance over all evaluation tasks[39]. Very recently, several attempts have been made to incorporate prompt-based learning to molecular design[40-43]. These works combine SMILES representation of molecules with other modalities including chemical structure texts[40, 41], pharmacological properties[41-43], medical description texts[42], and protein pocket[43]. In [43], Gao et al. propose a unified model called PrefixMol considering both chemical properties and binding pocket via generative pre-trained transformer (GPT). The pocket information is transformed into an embedding by geometric vector transformer (GVF) and used as a prefix condition together with other conditional embeddings. PrefixMol demonstrates excellent performance in single and multiple conditional molecular generation. But still, PrefixMol is an autoregressive model, and the global context of ligands are lost during the generation process. Moreover, their model treats pocket residues equally and the outputs are 1D SMILES representation, which makes it hard to apply in real scenarios.

## Discussion

In this work, we propose a novel diffusion model named InterDiff to guide the molecular generation from scratch by residue interaction prompt. Our model could generate molecules under certain interaction conditions with high statistical probability and better binding energies, which is critical for structure-based drug design. On the other hand, we demonstrate that InterDiff could be easily modified into a fragment-based generative model to help optimize molecules by introducing interactions with certain hotspots. The results show that InterDiff could generate new fragments that interact with hotspots based on a molecular scaffold. This characteristic is of wide insterests for modern in-silico drug design. Nevertheless, InterDiff still faces challenges such as the unsynthesizable structures presented in the generated molecules, which are also commonly seen in other methods. Although this may be the flaws in molecular reconstruction algorithms, efforts are needed to increase the molecular structural rationality. Potential solutions have been proposed as we discussed in the related work part. In the future work, we will attempt to optimize the sub-structures of generated molecules to ameliorate the drug-likeness and synthetic accessibility. Currently, the choice of the interaction prompts for residues mainly depends on the reference molecules or developer’s experience. Existing tools like FTMap may be helpful to solve this problem[44]. In addition, improving accuracy in describing π-π interactions could be important in future optimization.

## Methods

### Molecular diffusion model

We build our model upon the framework develop by Guan et al.[10]. Let ℳ = (*x, h*) denote the molecular 3D point cloud data with *x* = [*x*^(*L*)^, *x*^(*P*)^] *∈* ℛ^*N*×3^ and *h* = [*h*^(*L*)^, *h*^(*P*)^] *∈* ℛ^*N*×*M*^. In our setting, [*x*^(*L*)^, *x*^(*P*)^] indicate atom coordinates of ligand and protein, and [*h*^(*L*)^, *h*^(*P*)^] represent the atom categorial features, where *N* is the number of atoms. We use diffusion model to learn the distributions of protein-ligand complexes. Diffusion model learns two Markov processes, a diffusion process *q* and a denoising process *p*. Diffusion process adds Gaussian noise to data ℳ_*t*_ in time step *t*, where *t* = 0, *⋯, T* − 1 is the predefined time steps (*T* = 1000 in the implementation):

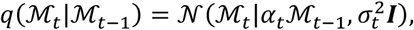

where *α*_*t*_ is the schedule that controls how much signals are preserved in the diffusion process and *σ*_*t*_ is the noise schedule that controls how much noises are added. For the 3D molecular point cloud data, the atom types are categorical data while the atom coordinates are continuous data. At time step *t*, We add Gaussian noise and uniform noise to the atom coordinate feature and atom type feature respectively[45]. Following the convention in [10, 45], the joint distribution states as:

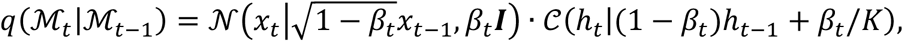

where *𝒞* indicates a categorical distribution with parameters after |, *β*_*t*_ is the variance schedules and *K* is the number of atom types with *k* = 1, *⋯, K*. In the implementation, the variance is reduced as the steps grow. For the generative denoising process, the posterior distribution *p*(·) can be computed in a close form by the Bayesian formula:

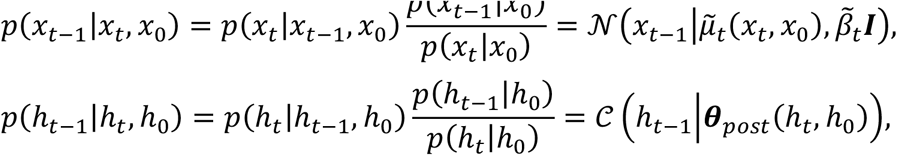

Where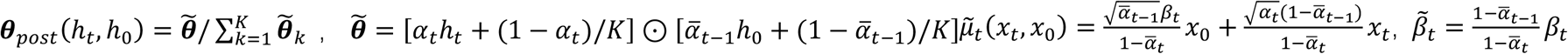 and *α*_t_=1−*β*_t_ 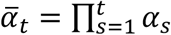. In the denoising process, *x*_0_ and *h*_0_ are approximated by neural network, and we denote the approximation of *x*_0_ and *h*_0_ as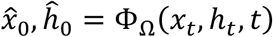, ĥ_0_ = Φ_Ω_(*x*_*t*_, *h*_*t*_, *t*), where Φ is a neural network parameterized by Ω. The training objective is the summation for atom coordinates and atom types. The atom coordinate loss states as:

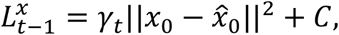

where *γ*_*t*_ is the weight for MSE loss and *C* is a constant. In the implementation, we set *γ*_*t*_ = 1 for all time steps. The atom type loss is computed by KL-divergence of two categorical distributions:

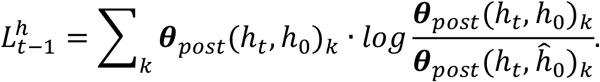

The final loss is calculated by the weighted summation of MSE loss, KL-divergence and a classification loss:

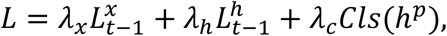

where *λ*_*x*_, *λ*_*h*_ and *λ*_*c*_ are the weight for MSE loss, KL-divergence and classification loss respectively and *h*^*p*^ indicates the protein atom features. The classification loss classified the protein atom features according to their atomic interaction types and the cross entropy loss are used in the experiment.

### Equivariant diffusion under prompt guidance

In this section, we elaborate our proposed InterDiff model. InterDiff is a graph neural network in which the atom denotes the nodes and the Euclidean distance between atoms denotes the edges. We define an edge among two nodes when the Euclidean distance is below 7 angstroms. Let 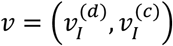denotes the interaction prompts, where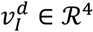 are one-hot representation prompts, *I ∈* (*cation* − *pi, halogen, hydrogen, pi* − *pi*) and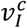 are learnable continuous embeddings. The atom node features are also encoded by one-hot vectors and transformed by a single linear layer: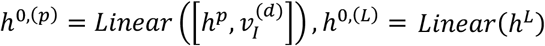.

InterDiff is composed of six equivariant block layers (Figure 6), and each block consists of three modules. Formally, the first module updates the node features:

**Figure 6.**
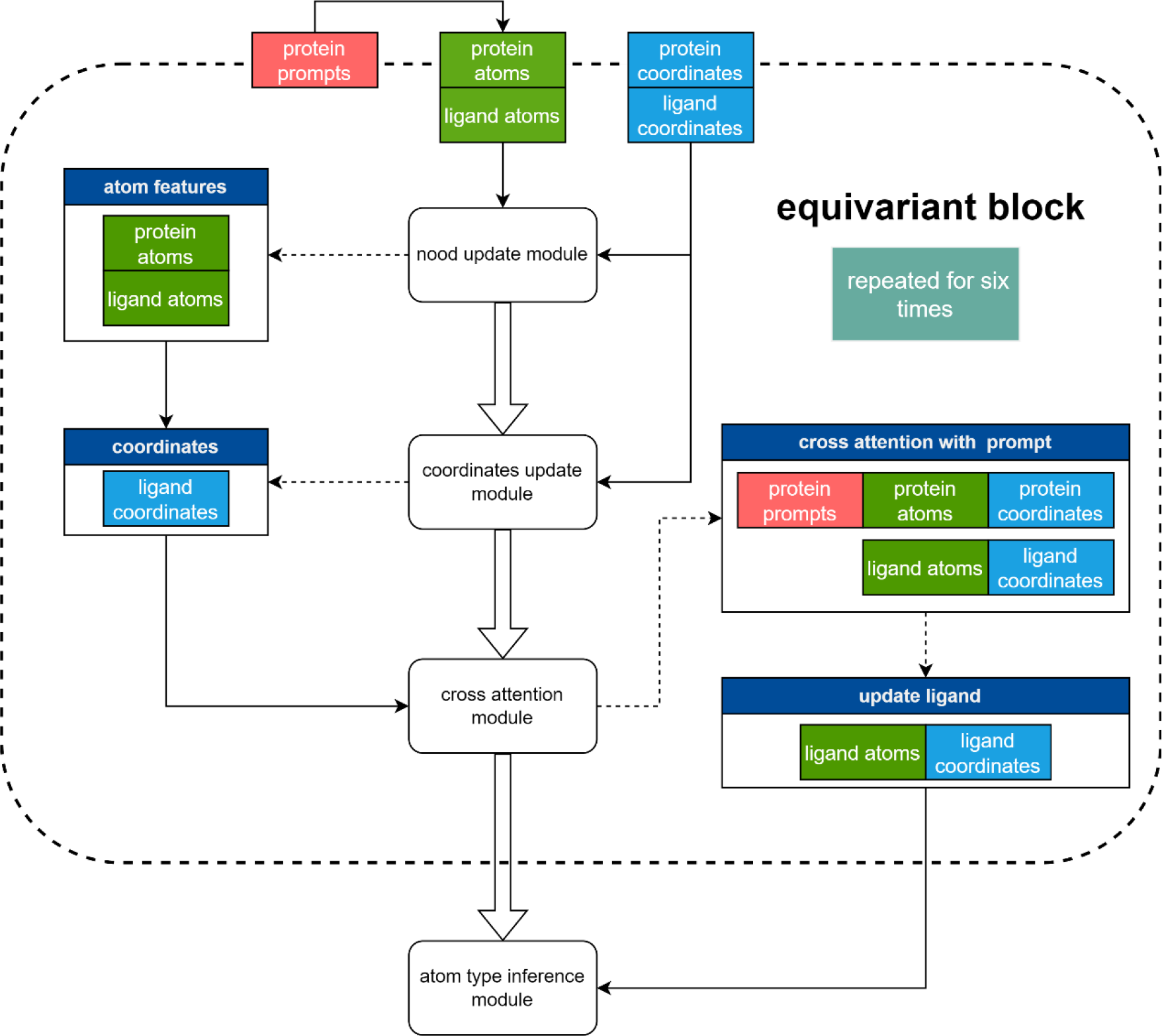
Structure of equivariant block used in InterDiff. Modules are represented by rounded rectangle with white context while data are shown by rectangle with distinct colors. Input and output flows are shown with arrowhead and dashed arrowhead respectively.

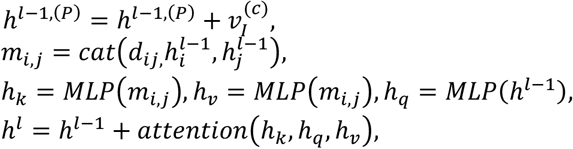

where *h*^*l*,(*P*)^ indicates protein atom features in *l* th layer, *d*_*ij*_ is the Euclidean distance between atom *i* and *j*, and *cat*(·) indicates the concatenation operation. The second module updates the ligand coordinates:

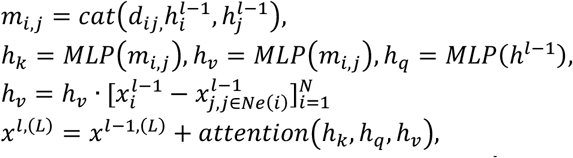

where *x*^*l*,(*L*)^ indicates ligand atom coordinates in *l*th layer,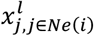 represents that *j*th atom coordinates and *j* is the neighbors of atom *i*. The third module updates ligand atom features and coordinates simultaneously with cross attention:

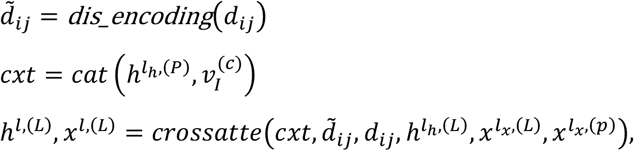

where *dis_encoding*(·) is the encoding of distance matrix between ligand and protein,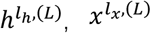and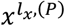 indicate the node features and coordinates of ligand and coordinates of protein from the first and second module. For the distance encoding, we use multilayer perceptron in the implementation. The *crossatte*(·) is computed as follows:

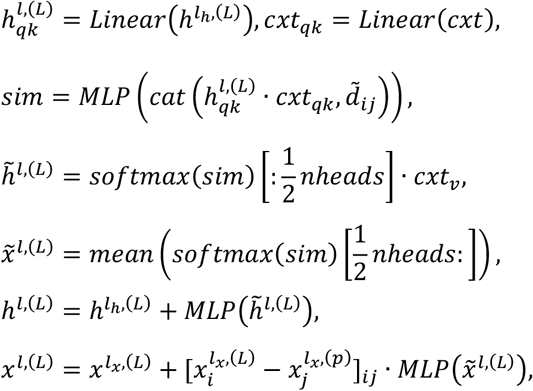

where *nheads* is the number of heads in cross attention mechanism, *sim* is the attention map between ligand and protein. Noted that we ignore the operations of dimension rearrangement in above formulas and simplify Einstein summation to dot product here. Identical to previous work [35, 46, 47], we use ‘subspace-trick’ by limiting the center of mass (CoM) of training samples to zero to ensure the model can achieve translation invariance in the generative process. For the SE(3)-equivariance of Markov transition, the proof of the first two modules is similar to [10] and we prove the equivariance for the cross attention module in the supplementary.

### Training and sampling details

InterDiff consists of 6 equivariant blocks and each block has three modules with transformer like structure. The diffusion steps are set to 1000 in training and sampling. We utilize a sigmoid *β* scheduler for atom coordinates and a cosine *β* scheduler for atom types. The number of heads is 16 for the first two modules and 32 for the cross attention module. The dimension is 128 for the atom features and interaction prompt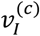. We use Adam[48] method to optimize the model with an initial learning rate 0.001, betas=(0.95,0.999) and the batch size is set to 8. A ‘plateau’ scheduler was applied to decay the learning rate with a factor 0.8 when the validation performance is stuck for 4 evaluation steps. The minimum learning rate is 1e-6. The loss weight is 100 for atom type loss and 1 for MSE loss. We train InterDiff on one NVIDIA V100S GPU and the model converges within 32 hours. In addition, we empirically found that the model could be further improved on validation set (randomly selected from training data for validation) when fix the prompt embedding and fine tune after convergence.

In the sampling process, the interaction prompts for each sample are provided in keeping with the molecule in test set. The center of mass is subtracted from the coordinates of protein atoms and the number of atoms is sample according to the pocket size (Figure S3). The initial coordinates of ligand atoms are sampled from a normal distribution and atom types are sampled from a Gumbel distribution and then transformed into one-hot vectors.

### Featurization of atoms and distance

Atoms in ligand and protein are represented by one-hot vector initially and then transformed by a linear layer. We use a mixed representation for protein atoms and ligand atoms as described in [10]. Specifically, the protein atom features encode the information about amino acid types, atom types and whether the atom is backbone atoms. The ligand atom features encode the atom types and aromatic information. The distance between atoms and bond types are used to construct graph edges. Four types of bonds are considered by one-hot vector, which indicates the connection between ligand atoms, protein atoms, ligand-protein atoms and protein-ligand atoms. The edge feature are then encoded by gaussian radial basis functions with learnable parameters of mean and variance, for the details please refer to [49].

### Characterizations and parameters of interactions

In this paper, we consider four types of interactions, and the characterizations of interactions are consistent with BINANA2. Cation-π interactions comprise of a charged functional group and an aromatic ring. The coordinate of charged functional groups is projected to the plane of the aromatic ring and cation-π interaction is accepted if the distance of two center points between pairs is less than a threshold. π-π interactions have two types of forms, pi-pi stacking (face to face) and T-stacking (edge to face). To detect π-π interactions, distance of the projection of center points on two aromatic rings and the angle of two vectors normal to planes for each ring are calculated. If the distance and angle satisfy certain thresholds, a π-π interaction is identified. Hydrogen bond is composed of a hydrogen bond donor and a hydrogen bond acceptor. In BINANA2, thiol, amine, and hydroxyl groups are allowed as donors and nitrogen, sulfur and oxygen atoms can act as receptors. Likewise the distance between donor and receptor and the dihedral angle between hydrogen atoms, donor and receptor must locate in a certain range. Halogen bonds also consist of a donor and a receptor. The donors include O-X, N-X, S-X, and C-X, where X is F, Cl, Br, or I. The acceptors could be nitrogen, sulfur and oxygen atoms. The threshold of distance for halogen bonds tends to be longer than hydrogen bonds and the dihedral angle is the same. The details of threshold values are listed in Table S2.

## Data availability

The CrossDocked 2020 can be obtained at https://bits.csb.pitt.edu/files/crossdock2020/; Structured models used in studies are deposited in Protein Data Band with accession codes 6oik, 3uon, 8azx and 8azv. The source code are available on Github: https://github.com/zephyrdhb/InterDiff.

