## Supplementary figures and tables for "Guided Diffusion for molecular generation with interaction prompt"

#### Additional figures and tables

| Ring | Pocket2Mol | AR | TargetDiff | InterDiff |
|---|---|---|---|---|
| 3 | 0.2% | 30.9% | 0.0% | 0.0% |
| 4 | 0.1% | 0.8% | 4.2% | 11.8% |
| 5 | 16.7% | 20.2% | 32.4% | 24.7% |
| 6 | 77.6% | 43.8% | 42.2% | 40.7% |
| 7 | 3.9% | 2.3% | 15.9% | 14.6% |
| 8 | 1.2% | 1.5% | 4.2% | 7.6% |
| 9 | 0.3% | 0.5% | 1.2% | 0.6% |

Table S1： Percentage of ring sizes in InterDiff and baseline methods.

| Parameter name | Value | Description |
|---|---|---|
| hydrogen_bond_dist_cutoff | 3.2 Å | Donor/acceptor distance cutoff for hydrogen bonds |
| hydrogen_halogen_bond_angle_cutoff | 40.0° | Donor/acceptor distance and angle cutoff, in degrees, for hydrogen and halogen bonds |
| halogen_bond_dist_cutoff | 5.5 Å | Donor/acceptor distance cutoff for halogen bonds |
| pi_pi_interacting_dist_cutoff | 4.4 Å | Ring-center distance cutoff for detecting pi-pi stacking and T-shaped interactions |
| pi_stacking_angle_tolerance | 30.0° | pi-pi stacking angle cutoff, in degrees |
| t_stacking_angle_tolerance | 30.0° | pi-pi T-shaped angle cutoff, in degrees |
| t_stacking_closest_dist_cutoff | 5.5 Å | Atom-atom distance cutoff for detecting pi-pi T-shaped interactions |
| cation_pi_dist_cutoff | 6.6 Å | Charged-moiety/ring-center distance cutoff for cation-pi interactions |

Table S2： Summary of parameters of BINANA2 in identifying four interactions.

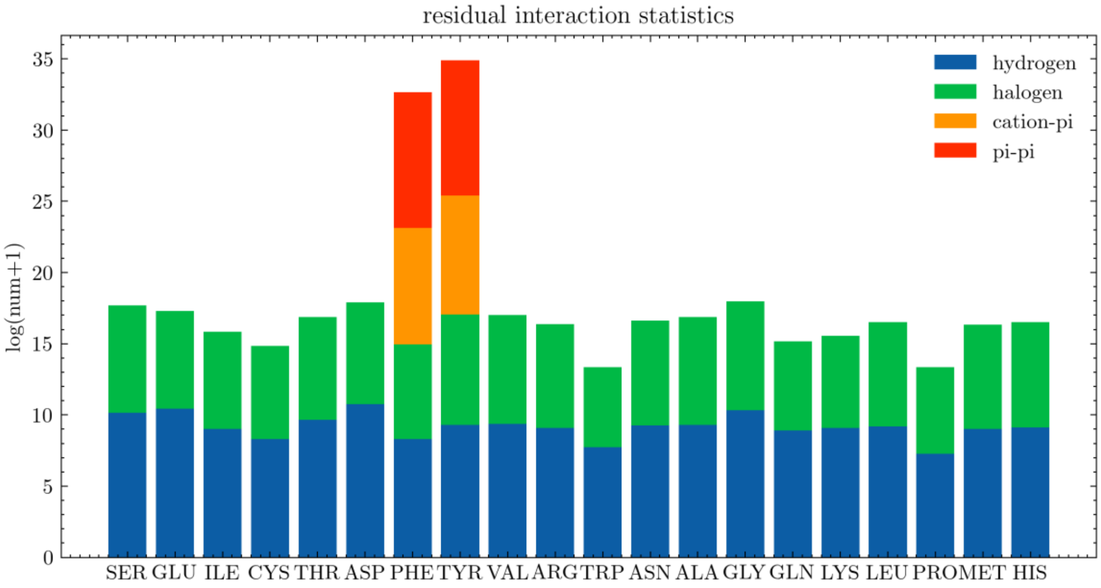


Figure S1: A bar plot of interactions in different protein residues. Y axis is illustrated with logarithmic scale to balance the height of the bar.

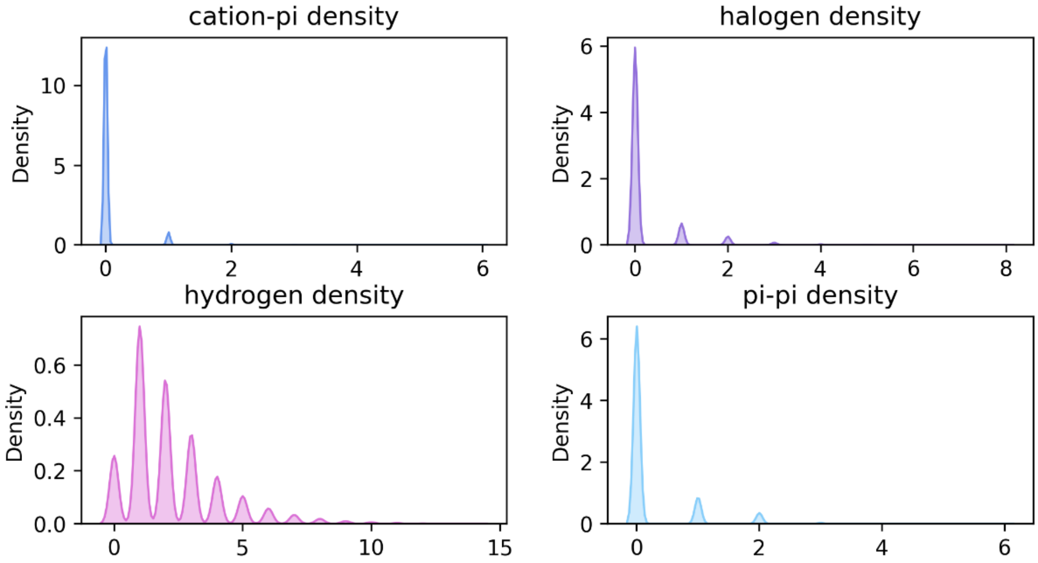


Figure S2: Density distributions of interactions in CrossDocked 2020 training set. The number of four interactions were count for each protein-ligand complex.

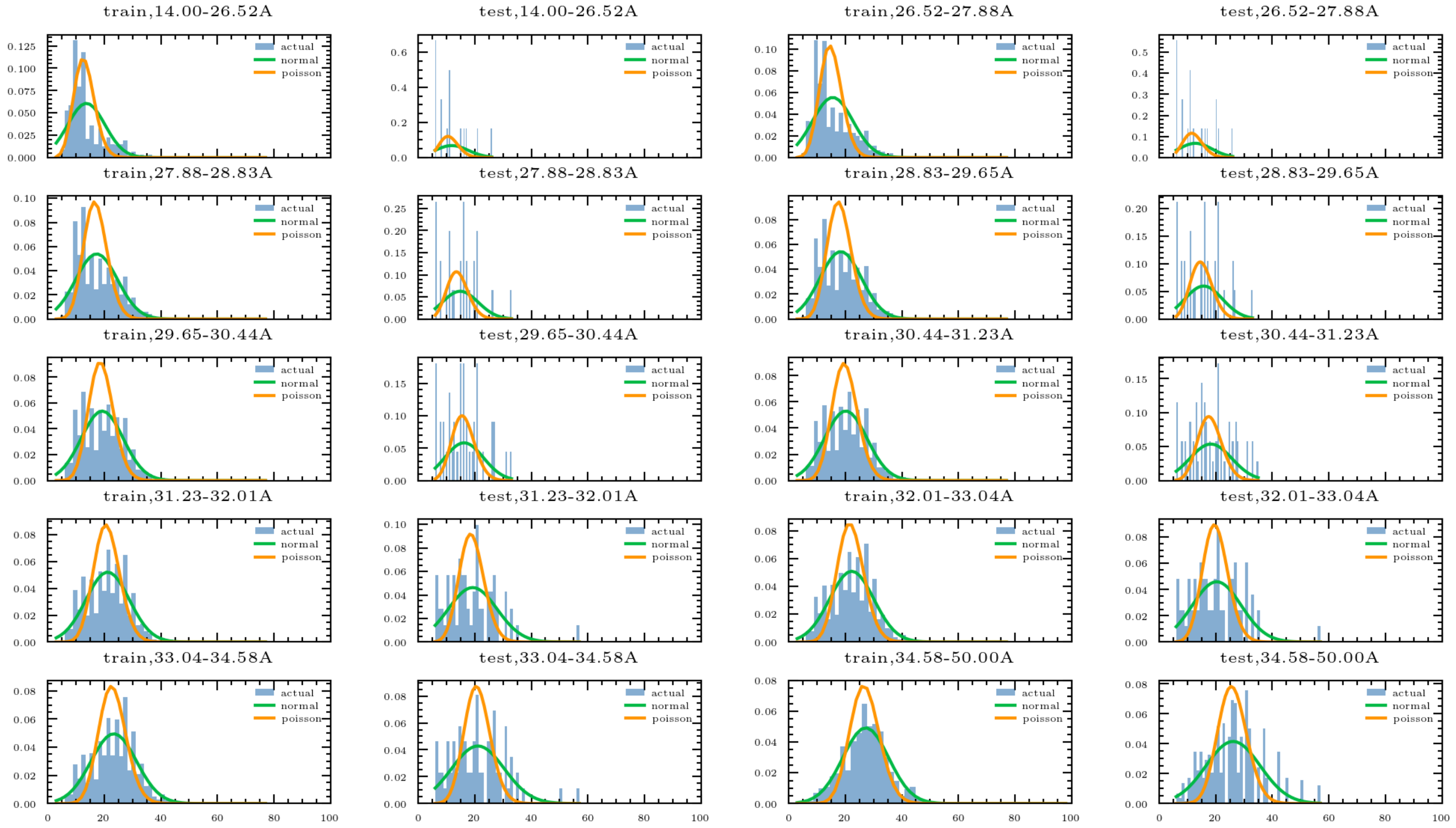


Figure S3: Distributions of number of ligand atoms in different pocket sizes. Pocket sizes are defined by the median distance of the 10 farthest of pocket atoms. We divide the pocket sizes range into 10 quantiles and plot the distributions of number of ligand atoms of each bin in training set and test set. There is a clear trend that the larger the pocket size is, the more the number of ligand atoms will have. In addition, we can see that the distribution is similar for the training and test set in each bin. The distribution in training set can be generalized to test set.

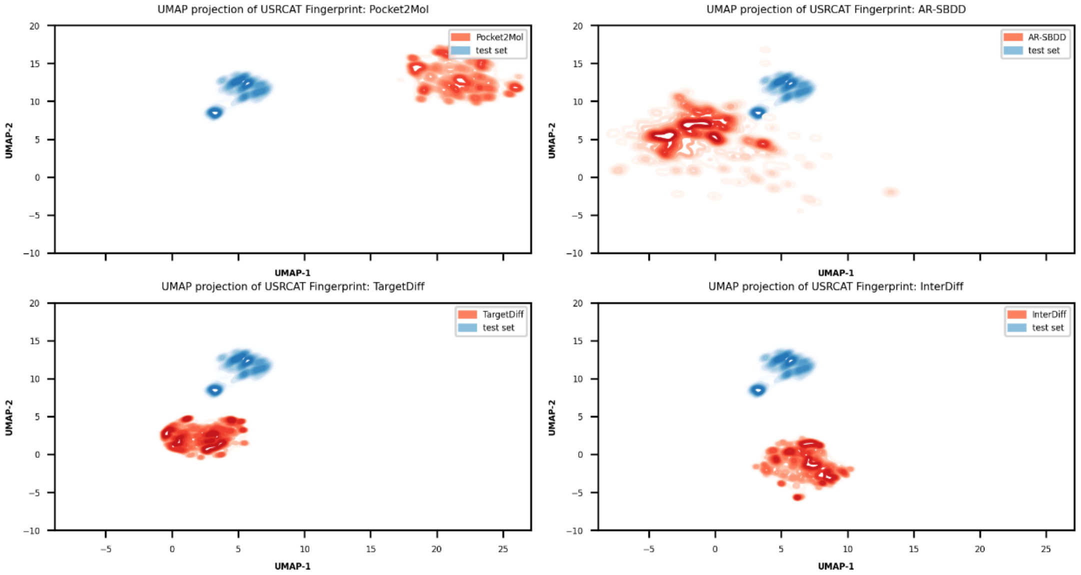


Figure S4: UMAP projection of USRCAT fingerprint in 2D space[1, 2]. The depth of color indicates the density value. The test set are colored with blue and the results from models are colored with red.

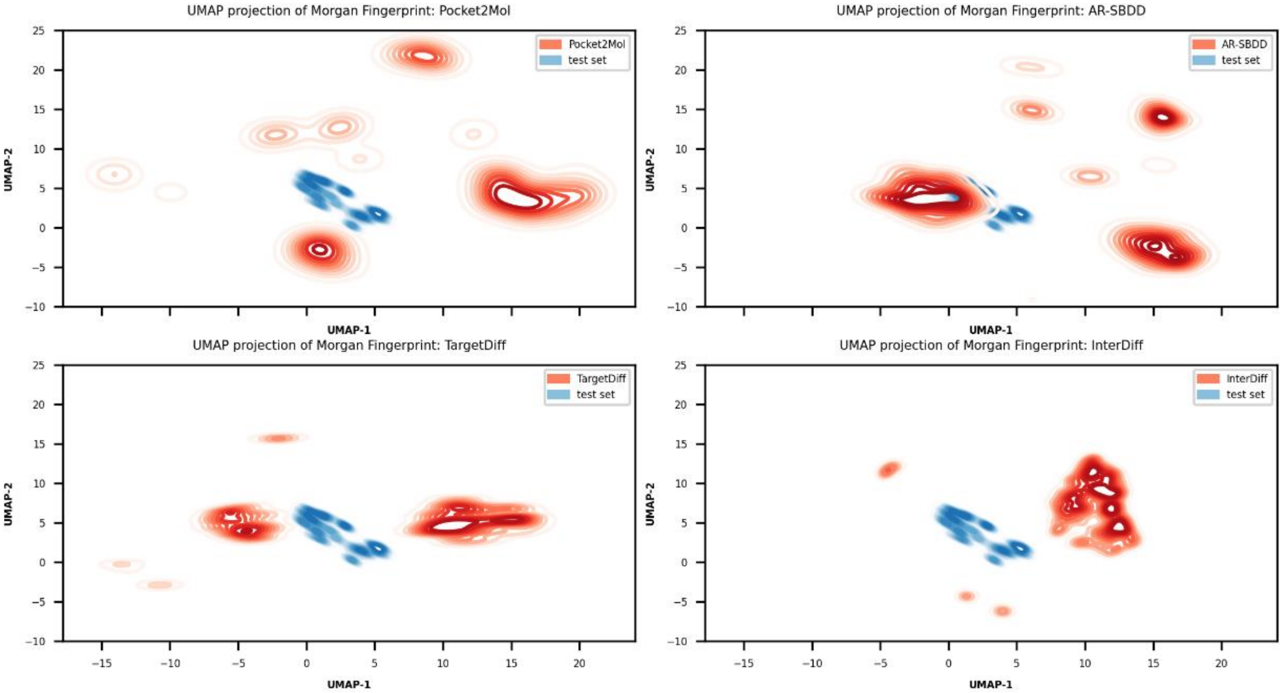


Figure S5: UMAP projection of Morgan fingerprint in 2D space[1]. The depth of color indicates the density value. The test set are colored with blue and the results from models are colored with red.

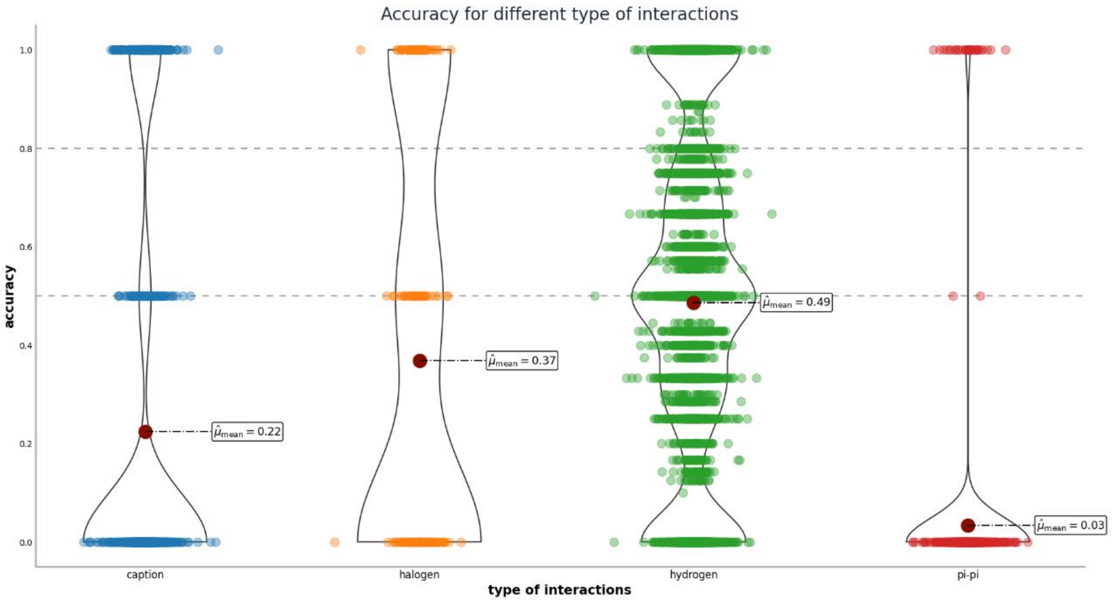


Figure S6: Accuracy of InterDiff in designing four types of interactions in test set.

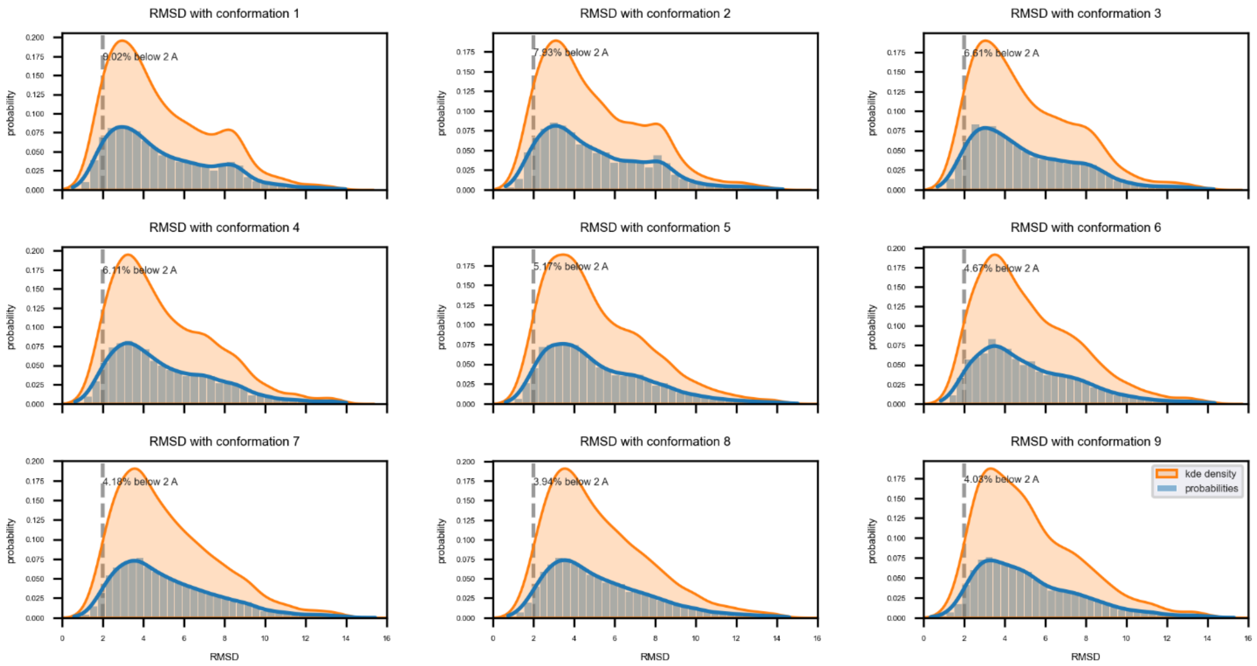


Figure S7: RMSD between original conformations generated by InterDiff and docked conformations in test set. 9 conformations are generated by QuickVina for each molecule and the percentage of RMSD below 2 angstrom is calculated.

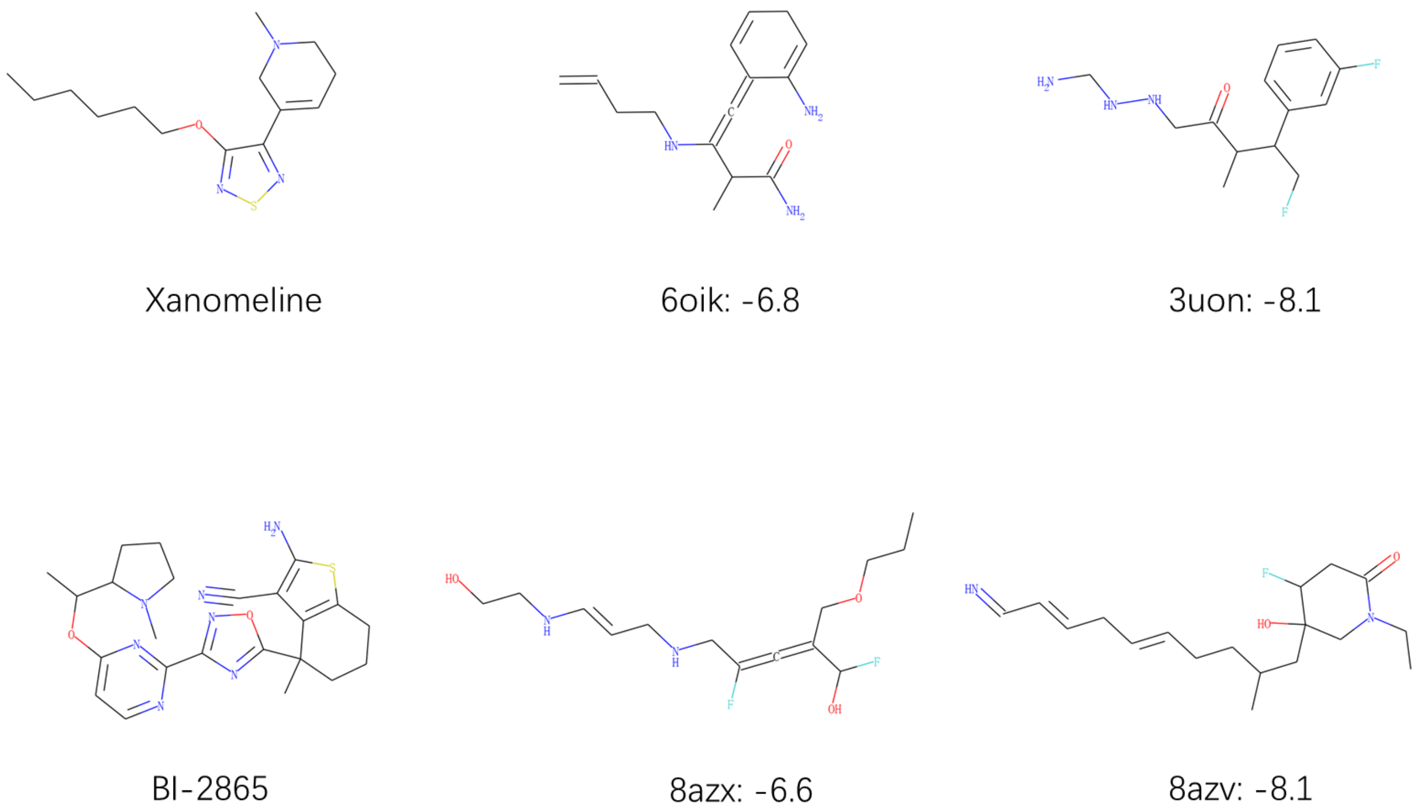


Figure S8: Native drugs and generated molecules for two protein targets, mAChR (top row) and KRAS (bottom row). The docking pose are illustrated in figure 4 and figure 5. Generated molecules are labeled with PDB accession codes and vina scores.

| PDB code | Target | interaction | Description |
|---|---|---|---|
| 6oik | mAChR active state | [(TRP,422,cation-π)] | The interactions are obtained by docking with QuickVina |
| 3uon | mAChR in-active state | [(TYR,104,cation-π),(TYR,403,cation-π)] | The interactions are obtained by docking with QuickVina |
| 8azx | KRAS mutant G12C | [(GLU,63,hydrogen),(ASP,69,hydrogen),(HIS,95,cation-π)] | The interactions are obtained from cocrystal structure |
| 8azv | KRAS wild type | [(GLU,63,hydrogen),(ASP,69,hydrogen),(TYR,64,cation-π),(HIS,95,cation-π)] | The interactions are obtained from cocrystal structure |

Tab S3: Interactions of targeting drugs in two protein targets.

### Proof of SE(3)-equivariance of cross attention module in generative Markov transition

The proof of SE(3)-equivariance in node and coordinate updating modules are the same as Guan's work[3], here we prove the equivariance of cross attention. Denoting the SE(3)-transformation as $T_g$, the equivariance of a parametrized module states as $m_\theta\left(T_g(\mathcal{M})\right) = T_g(m_\theta(\mathcal{M}))$ for a protein-ligand complex $\mathcal{M}$. Since the atom features are always SE(3)-invariant, we prove the atom coordinates update in cross attention module.

First, recall that the distance between atoms is encoded by $dis\_encoding(\cdot)$, which is invariant to SE(3) transformation:

$$\begin{aligned}\tilde{d}_{ij} &= dis\_encoding\left(||T_g(x_i) - T_g(x_j)||^2\right)\\ &= dis\_encoding\left(||(Rx_i + b) - (Rx_j + b)||^2\right)\\ &= dis\_encoding\left(||Rx_i - Rx_j||^2\right)\\ &= dis\_encoding\left(\left(x_i - x_j\right)^T R^T R\left(x_i - x_j\right)\right)\\ &= dis\_encoding(d_{ij}),\end{aligned}$$

where $R$ is a rotation matrix and $b$ is a translation vector. Similarly, the deviation, $\tilde{x}^{l,(L)} = mean\left(softmax(sim)[nheads:] \cdot d_{ij}\right)$ is also invariant to SE(3) transformation. Then, for update equation $x^{l,(L)} = x^{l_x,(L)} + [x_i^{l_x,(L)} - x_j^{l_p,(L)}]_{ij} \cdot Linear\left(\tilde{x}^{l,(L)}\right)$, denotes as $\phi_\theta(x^{l,(L)})$, we have:

$$\begin{aligned}\phi_\theta(T_g\left(x^{l,(L)}\right)) &= T_g\left(x^{l-c,(L)}\right) + [T_g(x_i^{l_x,(L)}) - T_g(x_j^{l_x,(p)})]_{ij} \cdot MLP\left(\tilde{x}^{l,(L)}\right),\\ &= Rx^{l_x,(L)} + b + [Rx_i^{l_x,(L)} - Rx_j^{l_x,(p)}]_{ij} \cdot MLP\left(\tilde{x}^{l,(L)}\right)\\ &= R(x^{l_x,(L)} + [x_i^{l_x,(L)} - x_j^{l_x,(p)}]_{ij} \cdot MLP\left(\tilde{x}^{l,(L)}\right)) + b\\ &= Rx^{l,(L)} + b\\ &= T_g\left(\phi_\theta\left(x^{l,(L)}\right)\right)\end{aligned}$$

So the cross attention is SE(3)-equivariant with respect to the input. By leveraging the conclusion from [4, 5] that a SE(3)-invariant initial density and a SE(3)-equivariant Markov transition function can guarantee an invariant likelihood with respect to $T_g$, we can draw a conclusion that our model is likelihood invariant to rotating and translating of protein-ligand complex.
